# Transcriptome Universal Single-isoform COntrol: A Framework for Evaluating Transcriptome reconstruction Quality

**DOI:** 10.1101/2025.08.23.671926

**Authors:** Tianyuan Liu, Alejandro Paniagua, Fabian Jetzinger, Luis Ferrández-Peral, Adam Frankish, Ana Conesa

## Abstract

Long-read sequencing (LRS) platforms, such as Oxford Nanopore and Pacific Biosciences, enable comprehensive transcriptome analysis but face challenges such as sequencing errors, sample quality variability, and library preparation biases. Current benchmarking approaches address these issues insufficiently: BUSCO assesses transcriptome completeness using conserved single-copy orthologs but can misinterpret alternative splicing as gene duplications, while spike-ins (SIRVs, ERCCs) oversimplify real-sample complexity, neglecting RNA degradation and RNA extraction artifacts, thus inflating performance metrics. Simulation algorithms are limited to recapitulate this complexity. To overcome these limitations, we introduce the Transcriptome Universal Single-isoform Control (TUSCO), a curated internal reference set of genes lacking alternative isoforms. TUSCO evaluates precision by identifying transcripts deviating from reference annotations and assesses sensitivity by verifying detection completeness in human and mouse samples. Masking TUSCO transcripts—and optionally inserting decoy splice variants—creates a ‘novel-isoform’ challenge that assesses recovery of the true, now-unannotated isoforms. Our validation demonstrates that TUSCO provides accurate and reliable benchmarking without external controls, significantly improving quality control standards for transcriptome reconstruction using LRS.

## Introduction

Advances in long-read sequencing (LRS) technologies, such as Oxford Nanopore Technologies (ONT) and Pacific Biosciences (PacBio), have revolutionized transcriptomic studies by enabling the comprehensive profiling of full-length RNA molecules^1^. These emerging LRS platforms allow precise quantification of isoform diversity and in-depth characterization of alternative splicing at bulk^2,3^ and single-cell^4,5^ levels, tasks that short-read methods struggle to accomplish^6^ (reviewed in ^7^). Despite these breakthroughs, the Long-read RNA-Seq Genome Annotation Assessment Project (LRGASP) Consortium and other benchmarking efforts^8–12^ have shown that transcript identification, particularly the accurate detection of novel isoforms absent from current annotations, remains challenging. Factors such as sample quality, library preparation, the RNA capture by the instrument, sequencing errors, as well as processing choices of the analysis method of choice may introduce biases and compromise the accuracy of transcript calls. This has motivated the development of tools and strategies for the quality evaluation of long-read transcriptomics data.

Comprehensive quality control (QC) is critical for identifying potential biases and technical artifacts in LRS data. Read-level QC tools, such as LongQC^13^, MinKnow (https://github.com/nanoporetech/minknow_api), SQANTI-reads^14^, and PycoQC (https://github.com/a-slide/pycoQC)^15^, assess sequencing quality using metrics including read counts, length distribution, basecalling accuracy, adapter contamination, GC content, and coverage, among others. At the transcript level, SQANTI3 is widely used for evaluating the structural novelty of transcripts, integrating complementary data such as short-read alignments, CAGE peaks, poly(A) motifs, and poly(A) sites, and machine learning to improve confidence and curation of long-read defined transcriptome^16^. Although these tools yield valuable insights into data quality, they do not offer an absolute ground truth and therefore cannot provide formal benchmarking capabilities.

Current long-read benchmarking frameworks have adopted and adapted existing short-read methods. For example, BUSCO, which measures completeness by comparing reconstructed transcriptomes against a database of conserved single-copy orthologs^17^, has been used by several LRS assessment studies^18–20^. However, BUSCO is limited in its capacity to evaluate isoform diversity, often misinterpreting alternatively spliced transcripts as gene duplications^21,22^. Another strategy is the utilization of spiked-in RNAs mimicking alternatively spliced transcripts, such as ERCCs^23^, SIRVs (lexogen.com/sirvs/) and Sequins^24^. These products offer LRS-tailored benchmarking capabilities but fail to capture the complexity of actual RNA samples, including RNA degradation patterns. As such, spike-ins tend to overestimate the performance of LRS methods. Finally, several data simulation algorithms including computational strategies to simulate novel transcripts have been proposed^25–27^. While these tools can generate comprehensive ground truth datasets for method evaluation, they face challenges in faithfully simulating the complexity of the different sources of biases present in the data, and their results need to be complemented with additional benchmarking solutions.

Given these limitations, we developed the Transcriptome Universal Single Isoform COntrol (TUSCO), an internal ground truth for LRS transcript identification that can evaluate both real sample and technology artifacts. TUSCO consists of a curated set of widely expressed genes characterized by highly conserved splice sites and the absence of evidence for alternative isoforms in the Recount3^28^ database. TUSCO offers several benchmarking advantages. First, it detects false positives by identifying TUSCO-mapped transcripts that deviate from established annotations, providing a measure of precision in transcript identification and revealing partial transcripts that may result from low sample quality. Second, it uncovers false negatives by leveraging the widespread expression of TUSCO transcripts across human or mouse samples, thereby assessing the sensitivity and completeness of transcript detection methods. Third, in its *novel* mode, TUSCO enables the evaluation of performance in detecting novel transcripts under conditions of incomplete annotation. Moreover, TUSCO provides direct and effortless assessment as it eliminates the need to manage spike-in reagents or synthetic data. Our development of TUSCO satisfies a requirement in the LRS transcriptomics field by providing an internal standard for quality control.

## Results

### Selection and Validation of TUSCO Genes for Transcript Benchmarking

We built TUSCO by enforcing cross-annotation single-isoform agreement and population evidence for splice/TSS invariance (Figure 2a). Candidate genes had to present one annotated isoform with identical exon–intron structure and strand across GENCODE and RefSeq (plus MANE Select for human)^29–31^. We then screened for not annotated alternative splicing using large Illumina-based splice junction compendia:

**Figure 1:**
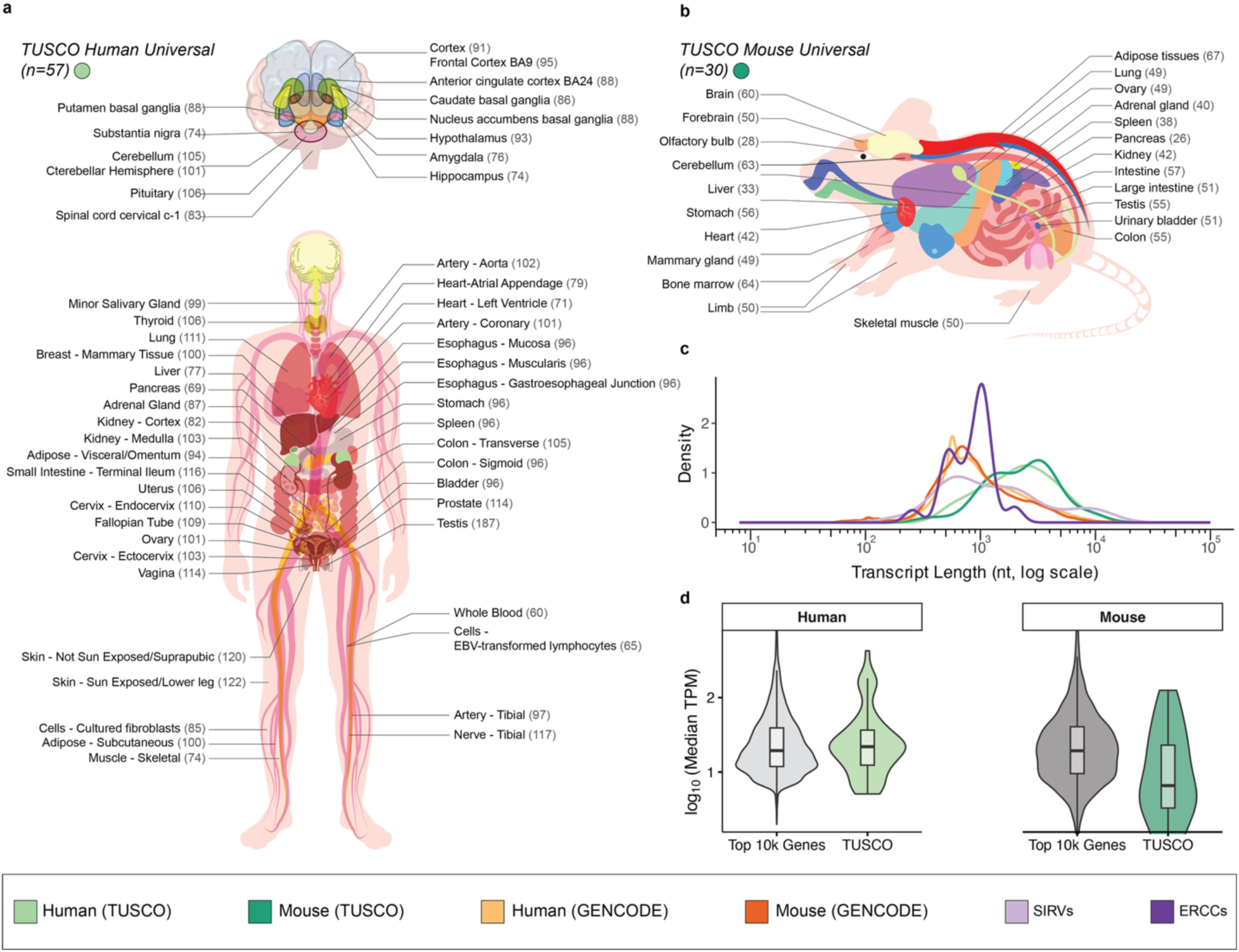
Comprehensive overview of TUSCO single-isoform gene sets in human and mouse across major tissues, comparing transcript length and expression levels against reference gene annotations. (a,b) Tissue representation for human (a) and mouse (b) TUSCO genes, showing the number of tissue-specific TUSCO sets identified across major organs. Each gene meets stringent single-isoform criteria—identical splice sites, transcription start sites, and transcription termination sites—verified across multiple annotation databases (GENCODE (v48, vM37), RefSeq (GCF_000001405.40, GCF_000001635.27), MANE Select (v1.4)). (c) Density plots (log10 scale) show the broad transcript length distribution of TUSCO transcripts (human/mouse) compared with all GENCODE (human/mouse) transcripts, SIRVs, and ERCCs. (d) Violin plots compare cross-tissue median expression (log10 TPM) of TUSCO genes (green) against the top 10,000 most highly expressed genes (gray) in human (left) and mouse (right) tissues based on the Bgee and GTEx source of gene expression. Embedded boxplots indicate the median (central line), interquartile range (box), and distribution range excluding outliers (whiskers).

**Figure 2:**
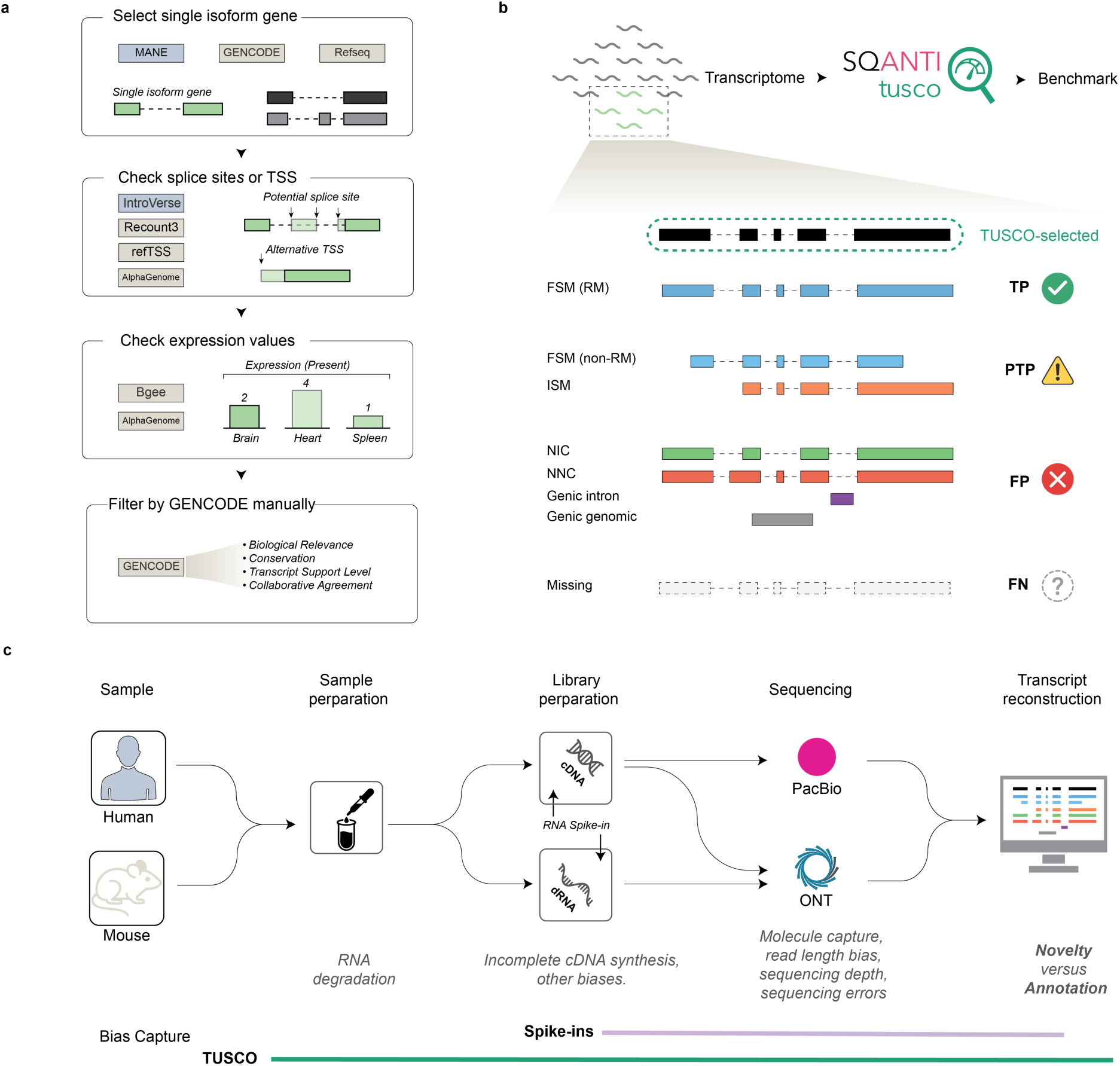
Overview of TUSCO gene selection and benchmarking framework. (a) Schematic illustration of the four-step pipeline for identifying single-isoform genes suitable for benchmarking. Candidate genes are cross-referenced across multiple annotation databases (MANE, RefSeq, and Ensembl for human; RefSeq and GENCODE for mouse), evaluated for potential alternative splice sites, and checked for broad, consistent expression in multiple tissues. (b) TUSCO leverages SQANTI3 structural categories to define benchmarking labels. True Positives (TP) are correctly identified TUSCO transcripts as Reference Match (RM); Partial True Positives (PTP) are those that align to alternative categories (FSM_non-RM or ISM). False Negatives (FN) indicate undetected TUSCO transcripts, whereas False Positives (FP) are transcripts with Novel in Catalog, Novel not in Catalog, Genic Intron, and Genic Genomic categories. (c) High-level workflow from sample preparation to computational analysis, highlighting how different biases (sample, protocol, platform, and transcript reconstruction) can affect user-defined transcriptome and different benchmarking approaches assess these biases.

IntroVerse (human)^32^ and recount3 junctions (human and mouse)^28^, removing loci with intragenic novel junction support above predefined thresholds (see Methods). TSS consistency was evaluated with refTSS (integrating FANTOM5, DBTSS, EPDnew, ENCODE)^33^; in human we applied both exon-overlap and ±300 bp CAGE-window checks, and in mouse we required the exon-overlap rule for single-exon genes (Methods).

Expression breadth was defined using Bgee present/absent calls with stringent prevalence cutoffs (human ≥99% of retained tissues; mouse ≥90%), reinforced by housekeeping genes from HRT Atlas^34,35^; tissue panels were derived as described in Methods. To further favor widely expressed, splice-invariant loci, we applied AlphaGenome filters on two metrics—Expression (tissue-median RPKM) and Splicing (maximum novel-splicing score/ratio)—using fixed species-specific cutoffs^36^. Remaining edge cases underwent manual review by GENCODE curators.

This process yielded tissue-resolved TUSCO gene sets (human: 65–187 per tissue; mouse: 28–67) and universal cores (human n = 57; mouse n = 30) consistently expressed across all tested tissues (Figure 1a,b). Compared with reference annotations and spike-ins, TUSCO spans endogenous size/structure ranges relevant for full-length detection: transcripts are comparable to RefSeq and longer than SIRVs/ERCCs; sets include both single- and multi-exon loci (Figure 1c; Figure S1). TUSCO genes also show expression distributions similar to highly expressed genes across tissues (Figure 1d) and, by AlphaGenome, higher Expression and lower novel-splicing scores than other single-isoform genes (Figure S2). Together, these properties support TUSCO as an endogenous, broadly expressed, splice-invariant ground truth for benchmarking transcript identification and completeness in long-read data.

### TUSCO-SQANTI3: A Unified Framework for Transcript Benchmarking

TUSCO leverages the SQANTI3 framework to assign LRS transcript models to benchmarking labels (Figure 2b). Hence, True Positives (TP) are TUSCO transcripts correctly identified by SQANTI3 as Reference Match (RM), i.e. transcript models matching a reference transcript at all junctions, and 5′ and 3′ ends within 50 nt from the annotated TSS or TTS, respectively. Partial True Positives (PTP) include transcripts classified as non-RM full-splice match (FSM) or incomplete-splice matches (ISM), implying significant deviations from the reference start or end positions, indicating RNA degradation or incomplete sequencing. False Negatives (FN) represent TUSCO transcripts that remain undetected at the gene level, possibly reflecting insufficient sequencing depth or biases in RNA capture. False Positives (FP) are transcripts mapped to TUSCO genes as Novel in Catalog (NIC), Novel not in Catalog (NNC), Genic Intron, Genic Genomic, or Fusion events, which may arise from sequencing artifacts, wrong transcript reconstruction or mapping errors.

These benchmarking labels allow for the calculation of standard performance metrics—Sensitivity, Precision, and Redundancy, which are provided in a dedicated TUSCO report. Seamlessly integrated with SQANTI3, TUSCO operates without requiring additional user input, enabling users to execute TUSCO analyses directly after completing SQANTI3 processing. The incorporation of the TUSCO report in the SQANTI3 results in an enhanced, comprehensive and unified quality control resource that both describes the properties of the long-reads data but also provides rigorous and standard performance metrics. Additionally, TUSCO enables a more complete assessment of the RNA sequencing workflow and capturing biases at every stage—from sample preparation to RNA extraction, library preparation, sequencing, and transcript reconstruction—unlike synthetic spike-ins such as SIRVs, which only gauge library preparation, sequencing, and transcript reconstruction (Figure 2c).

### Validation and Comparison between TUSCO and SIRVs

To assess TUSCO as an endogenous benchmark, we first compared its performance with respect to SIRVs using comprehensive data sets from the LRGASP project. Three major library preparation workflows— direct RNA (dRNA) ONT, cDNA ONT, and cDNA PacBio—were applied to both human WTC11 iPSC and mouse ES cells. Transcript models were generated using the FLAIR pipeline and evaluated against either TUSCO or SIRVs (Figure 3a).

**Figure 3:**
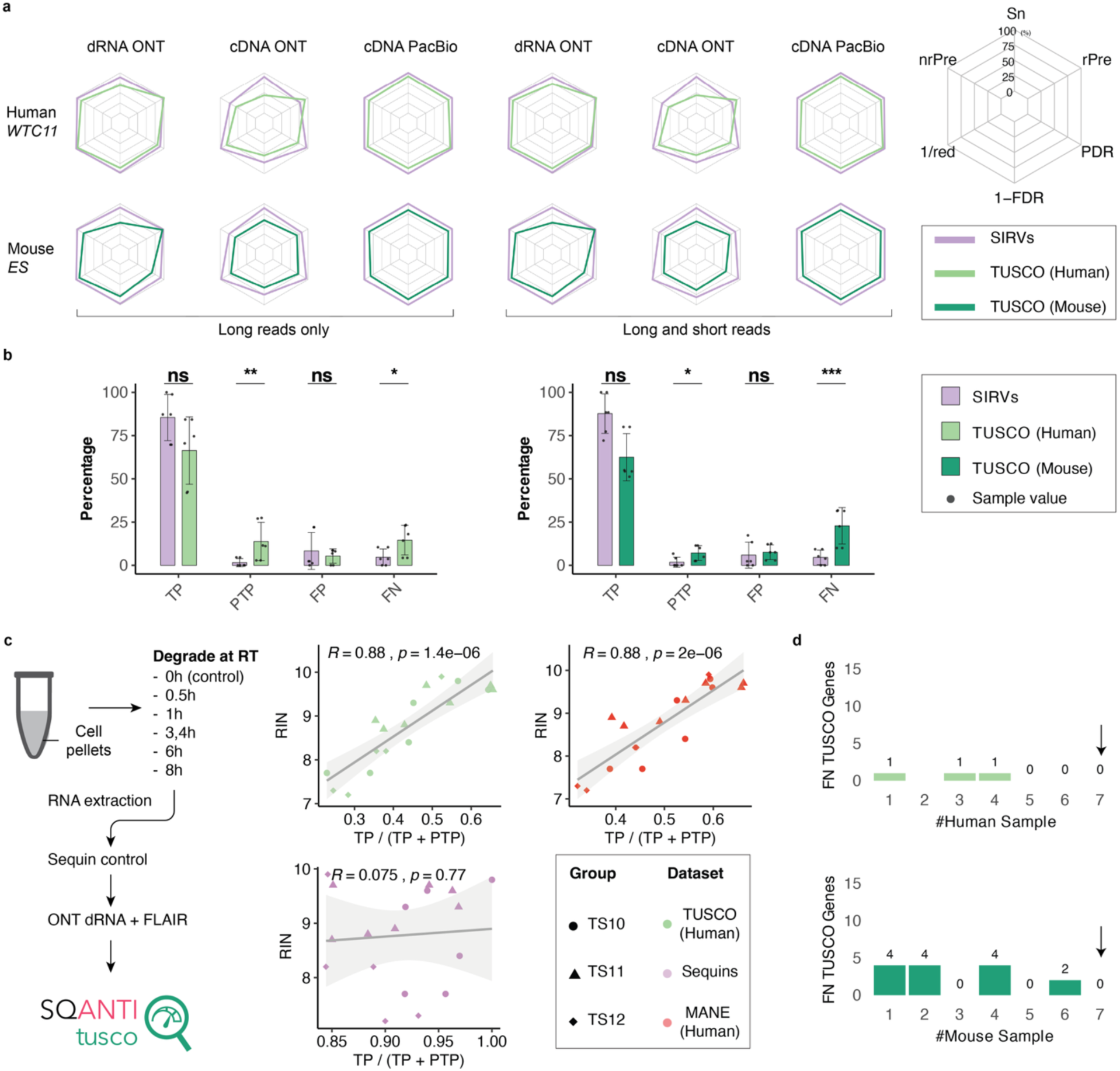
Comprehensive performance assessment of TUSCO and SIRVs for isoform detection across multiple sequencing platforms and pipelines, including the impact of RNA degradation on transcript recovery and analysis of false-negative genes. (a) Radar plots of sensitivity (Sn), precision (Pr), and redundancy (rPre) for human WTC11 (top row) and mouse ES (bottom row) samples processed with direct RNA (dRNA) ONT, cDNA ONT, or cDNA PacBio library preparation. Metrics were computed using either TUSCO (green and teal lines for human and mouse, respectively) or SIRVs (purple) (b) Bar charts comparing the frequencies (percentage) of true positives (TP), partial true positives (PTP), false positives (FP), and false negatives (FN) under TUSCO or SIRVs, shown for long-read-only pipelines (left) or combined long–short read pipelines (right). Significance between TUSCO and SIRVs is indicated by asterisks (ns = not significant; *p<0.05; **p<0.01; ***p<0.001). (c) Influence of RNA degradation on TUSCO and Sequin detection. A schematic (left) illustrates the experimental design, where RNA is degraded at room temperature (RT) for increasing times before sequencing. Scatter plots (right) show the correlation between RIN and the fraction of fully recovered transcripts for TUSCO (top) or Sequin (bottom). The fitted lines highlight the distinct relationships for each dataset (R² and p values indicated). (d) Identification of false-negative (FN) TUSCO genes across transcript reconstruction pipelines (top: human WTC11; bottom: mouse ES). For a total of 17 (human) and 20 (mouse) TUSCO genes not identified in at least one dataset, bar plots depict the number of FN TUSCO genes (y-axis) present in exactly k other datasets (x-axis) corresponding to six long-read protocols and one Illumina short-read dataset. Genes are labeled FN if they lack full-splice or incomplete-splice matches in long-read data or show no expression (TPM = 0) in short-read data. All FN were detected in at least one other dataset.

Across all library types and samples, TUSCO-derived metrics closely mirrored those obtained with SIRVs in sensitivity, precision, and redundancy (Figure 3a). To quantify this concordance, we calculated cosine similarity (cosim) scores between the two benchmarking approaches (Table S1). Values approaching 1 indicate strong agreement in evaluating transcript reconstruction performance but do not necessarily reflect absolute performance quality. The cDNA PacBio libraries, which were the best performing experimental option according to LRGASP results, consistently achieved cosim values near unity in both human and mouse samples, demonstrating that TUSCO and SIRV benchmarks equally recognize the performance of the high-quality data. Likewise, cDNA ONT, dRNA ONT, and hybrid long–short read pipelines (denoted “LS”) also exhibited high agreement (cosim: 0.97–0.99), reinforcing TUSCO’s utility as a reliable internal benchmark comparable to established spike-in controls, but also revealing small differences between the two QC datasets.

### TUSCO More Stringently Reflects Sample Preparation and Sequencing Depth

To understand these differences, we performed a detailed analysis of TUSCO genes across primary performance indicators. We compared the frequencies of true positives (TP), partial true positives (PTP), false positives (FP), and false negatives (FN) under TUSCO or SIRVs, for human and mouse datasets (Figure 3b). We found no significant differences between TUSCO and SIRVs in terms of TP and FP, confirming our previous conclusions when comparing performance metrics. To verify that TUSCO-identified FPs truly represent errors rather than alternative isoforms, we manually inspected these transcripts, revealing sequencing, mapping, or reconstruction errors unsupported by either long- or short-read data (Supplementary Material). In contrast, partial true positives (PTP) and false negatives (FN) differed significantly (one-sided paired t-test: human PTP p = 0.0098, FN p = 0.0125; mouse PTP p = 0.0212, FN p = 6.37 × 10^−4^) (Figure 3b), with both metrics higher under TUSCO. We hypothesize that because TUSCO represents sample endogenous transcripts, it is more sensitive to partial degradation or incomplete capture issues in sample quality and extraction that synthetic spike-ins like SIRVs or Sequins cannot fully replicate. To test this hypothesis, we applied TUSCO benchmarking to a set of ONT cDNA samples spanning a broad range of RNA integrity (RIN) values (Figure 3c)^44^. TUSCO assessment revealed a higher number PTP at lower RINs, reflecting the enhanced RNA degradation associated with low RIN samples. Notably, the fraction of fully recovered TUSCO transcripts—calculated as TP / (TP + PTP)—strongly correlated with the RIN value (R = 0.88, p = 1.410^−6^), whereas Sequins showed no correlation (R = 0.075, p = 0.77). Accordingly, TP/(TP+PTP) values separate clearly between benchmarks: TUSCO declines at low RIN and spans a wider range, while Sequins stay consistently high across the RIN spectrum. This near-ceiling behavior arises because spike-ins are intact and insulated from extraction and degradation, so they fail to expose partial-transcript artifacts that TUSCO detects.

We further examined false negative (FN) TUSCO genes across diverse assembly pipelines and sequencing technologies within the LRGASP dataset. We asked if we could find evidence of expression for all TUSCO genes in any of the LRGASP biological samples. We calculated the number of datasets where TUSCO genes were detected by at least one mapping read. We found that all TUSCO genes were detected in at least one dataset (Figure 3d), confirming their expression in the biological samples and their true False Negative status when not detected. Subsequently, we investigated the impact of sequencing depth and read-length quality on the occurrence of false negatives (Figure S2). Specifically, we correlated the number of false negatives with a composite metric defined as the logarithm of the total read count multiplied by the median read length. This analysis was conducted on pipelines utilizing exclusively long-read data as well as on those integrating both long-read and short-read sequencing. A pronounced negative correlation (Pearson r = –0.82, p = 1.1 × 10^−3^) was observed, indicating that as sequencing coverage and read length increase, the number of false negatives decreases. These findings suggest that the absence of TUSCO transcripts in specific pipelines predominantly reflects technical or coverage-based limitations, rather than a genuine lack of transcript expression, further validating the TUSCO dataset for benchmarking purposes.

We concluded that, by reflecting real sample pitfalls—including RNA degradation, library preparation biases, and variable read depth—TUSCO provides a realistic, endogenously anchored benchmark strategy that surpasses synthetic controls such as SIRVs and Sequins.

### TUSCO-novel Evaluates the Capacity of Reconstruction Tools to Detect Unannotated Isoforms

Since the TUSCO genes are well-annotated, they assess the capacity of long-read methods to detect known transcripts but do not reveal their accuracy in detecting novel transcripts, which often correspond to alternative isoforms of annotated genes^45–47^. To address this shortcoming, we developed a strategy to use the TUSCO framework to assess novel transcript calls. For each single-isoform TUSCO gene, we modified its transcript model in the reference annotation by removing one of the annotated splice junctions, inserting a plausible but artificial junction, and supplying this altered transcript model as the sole reference. This effectively updates the annotation of the single-isoform genes to reflect a non-existing transcript while omitting the real, expressed transcript (Figure 4a). Isoform reconstruction tools with capacity for detecting novel transcripts should call the true isoform despite a misleading annotation. We applied TUSCO-novel to Bambu, StringTie2, FLAIR, and the Iso-Seq + SQANTI3 ML pipeline across PacBio and ONT CapTrap and cDNA libraries in human and mouse (LRGASP). Performance with the native reference annotation (solid polygons, Figure 4b) was contrasted with performance under TUSCO-novel (dashed polygons).

**Figure 4.**
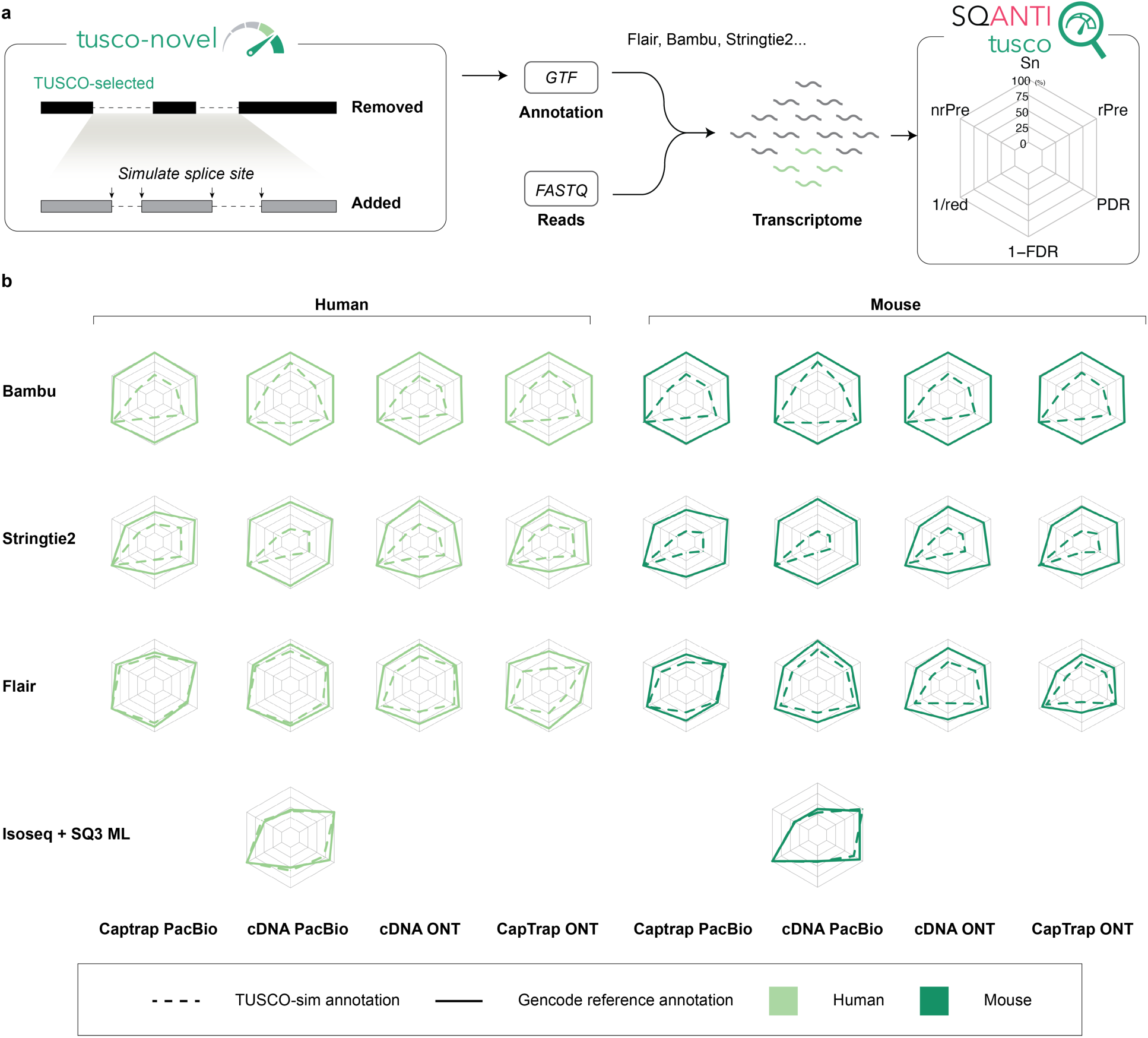
TUSCO-novel benchmark for evaluating multi-exon transcript discovery. **a** Schematic of the TUSCO-novel approach. The native TUSCO multi-exon isoform is removed from the annotation and replaced with a simulated transcript model carrying artificial donor and acceptor splice sites. The modified annotation is used as input to reconstruction pipelines to assess their ability to recover the true (but now unannotated) isoform. **b** Radar plots comparing performance metrics (sensitivity, precision, 1/FDR, redundancy, PDR, and 1/redundancy) for four reconstruction tools—Bambu, StringTie2, FLAIR, and Isoseq+SQ3 ML—across human and mouse samples using different library types. Solid lines represent results using the original reference annotation; dashed lines represent performance using the TUSCO- novel simulated annotation.

The results reveal clear distinctions in tool behavior. Bambu and StringTie2, which rely heavily on reference guidance, exhibited high performance when the true annotation was available but failed almost completely to recover the true TUSCO isoforms when only the novel simulation was provided. This sharp drop indicates that reference-driven tools struggle to detect truly novel splice junctions, even when read support is present (Figure 4b, Figure S3a-d). FLAIR demonstrated a more balanced profile, partially recovering the original TUSCO isoforms in the simulated novel context. However, the number of FPs and FNs remains elevated (Figure 4b, Figure S4e, f) under TUSCO-novel. Iso-Seq and SQ3 ML controlled FPs best across all pipelines, with 0 FPs reported under TUSCO-novel (Figure 43g, h). We hypothesize that this is because this pipeline combines the ability of Iso-Seq -reference-free data-driven method- for transcript discovery with the SQ3 filter that removes transcripts models lacking junction support from complementary data, thereby improving precision for novel transcripts. The relatively lower sensitivity and non-redundant precision reflect the lack of TSS or TTS adjustment against the reference of the Iso-Seq pipeline, which results in non-RM transcripts classified as partial true positives (Figure 4, Figure S4g, h). To verify this hypothesis, we calculated the read overlap over the true TUSCO transcript for PTP and TP separately in human and mouse. We found that, in human, TP have long-read coverage of 96.92 ± 10.22% of the TUSCO transcript length, dropping to 64.77 ± 62.90% for PTP; in mouse, TP coverage is 96.46 ± 9.72% versus 75.58 ± 29.56% for PTP. These results confirm the impact of read length on TUSCO transcript detection for reference-free methods (Figure S5).

Collectively, these findings demonstrate that TUSCO-novel provides a realistic, controlled framework to evaluate novel isoform discovery under practical conditions where the available reference may be misleading or incomplete. By simulating biologically plausible isoforms at real genomic loci, TUSCO-novel distinguishes between tools that strong base transcript calls on the annotation and those capable of reference-agnostic, data-driven transcript reconstruction—a key requirement for transcriptomics in emerging models, under-characterized tissues, or disease contexts.

### TUSCO helps assess replication in PacBio long-read transcriptomics

One of the most pressing questions in long-read RNA sequencing is how much replication is needed for consistent results. We asked if TUSCO could be used to evaluate how replication impacts sensitivity and precision. Five cDNA libraries were generated from mouse brain and kidney and sequenced on the PacBio Sequel IIe (∼5 M reads per sample; Supplementary Table 2). We computed TUSCO metrics in a multi- sample intersection mode—i.e., on transcripts consistently identified using FLAIR across the selected replicates (Fig. 5a).

**Figure 5.**
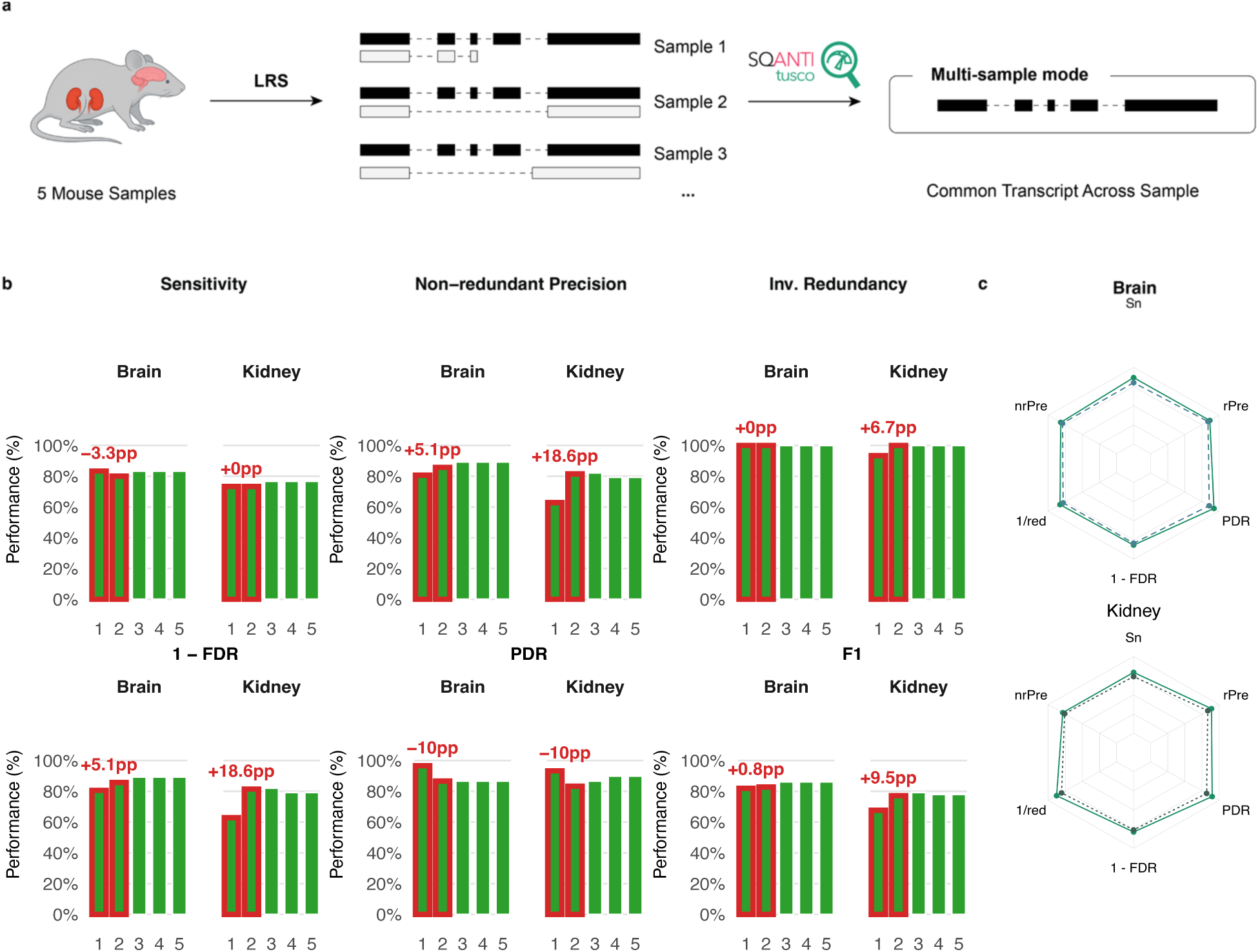
Replication gains and agreement between universal and tissue-specific TUSCO panels in mouse PacBio cDNA data. **a,** multi-sample (“intersection”) evaluation: five biological replicates per tissue (brain, kidney) were sequenced; SQANTI3 assigns transcript models; TUSCO metrics are computed on the transcript set common to the selected replicates. **b,** Performance versus replication (1→5 replicates) for brain and kidney. Bars show Sensitivity (Sn), non-redundant Precision (nrPre), inverse Redundancy (1/red), 1–FDR, PDR, and redundant Precision (rPre). Red callouts mark percentage-point changes relative to one replicate; note the large FDR reduction from one to two replicates, especially in kidney. **c,** Benchmark agreement between the universal mouse set (|U| = 30) and tissue-specific panels built with the same single-isoform criteria but requiring only tissue-consistent expression (brain: 60 genes; kidney: 42 genes). Values shown in the radar plots are computed per sample and then averaged across biological replicates within each tissue and panel type.

A single 5 M-read replicate already recovered most TUSCO transcripts (brain: Sn = 83.3%, PDR = 96.7%; kidney: Sn = 78.3%, PDR = 93.3%), but precision was lower (brain: Pr = 80.6%; kidney: Pr = 62.9%), leading to high False Discovery Rates (brain: FDR = 19.4%; kidney: FDR = 37.1%) Requiring transcript to be detected by ≥2 replicates sharply reduced false positives with minimal sensitivity loss. Precision increased to 85.7% (brain) and 81.5% (kidney) with FDR dropping to 14.3% and 18.5%, respectively. Also, Redundancy improved to nearly null in both tissues with 2 replicates. Sensitivity in the brain samples decreased slightly to 80.0% while this parameter was unchanged for kidney and the Positive Detection Rates dropped by 10 perceptual points in both datasets. F1 score was 81% for brain and 79% for the kidney samples. Interestingly, adding 3–5 replicates to the consistent detection did not modify significantly any of the performance metrics (Fig. 5b), with precision in brain stabilizing around ∼89.3% and sensitivity at ∼83.3%. For kidney sensitivity went to 76.7% and precision plateaued near 80% (Fig. 5b). These results reveal that TUSCO can be used to evaluate the effect of replication in transcript detection accuracy and suggest that, at least for the conditions of this experiment (mouse brain profiled by PacBio cDNA sequencing and processed with FLAIR) an optimal experimental setting for transcript detection would require at least 2 replicates to control the detection of spurious transcript models.

### TUSCO universal and tissue-specific sets give similar results

Previous analyses were conducted with the universal set of TUSCO genes, which is rather small (|U| = 30). We built tissue-specific TUSCO sets to expand loci and tested whether conclusions persisted. Tissue sets were constructed with the same single-isoform and invariance filters, requiring only consistent expression in the focal tissue. Using the same data, the brain (60 genes) and kidney (42 genes) panels produced benchmarking figures that were minimally lower than those of the universal set across in all cases; with cosine similarity between universal and tissue results was 0.9998 for brain and 0.9997 for Kidney (Fig. 5c). We concluded that the universal TUSCO set is a suitable benchmarking option for human and mouse samples, while tissue-specific sets provide a slighter higher coverage to fine-tune results.

## Discussion

The reconstruction of full-length transcriptomes using long-read sequencing technologies has revolutionized our understanding of transcript diversity. Yet, this advancement brings with it a pressing need for accurate and realistic benchmarking strategies. Our study demonstrates that synthetic spike-in controls, while useful, present significant limitations when used as stand-alone benchmarks. Tools like SIRVs and Sequins offer a controlled environment to assess transcript identification, but they fail to replicate key biological and technical variables that are intrinsic to real samples—such as RNA degradation, extraction bias, and natural variability across tissues. These shortcomings may result in artificially inflated performance estimates and obscure technology-specific limitations that are critical in challenging experimental contexts, such as clinical or disease-focused transcriptomic studies.

To address these gaps, we developed TUSCO, a benchmarking framework built around a highly curated set of single-isoform endogenous genes. These genes are robustly and consistently expressed across tissues, offering a more biologically relevant and nuanced ground truth to evaluate transcriptome reconstruction performance. TUSCO surpasses synthetic benchmarks by reflecting the same degradation, extraction biases, and length distribution characteristics of native RNAs, particularly in the medium-to-long transcript length range, which is underrepresented in many synthetic datasets. This results in a more representative assessment of sequencing performance.

Moreover, because TUSCO genes are endogenous and tissue-resolved, they enable benchmarking across a wide variety of experimental settings and sample qualities. This versatility makes TUSCO particularly valuable for several practical applications. For example, we show that the framework can inform trade-offs between sequencing depth and number of replicates, a common dilemma in experimental design. TUSCO also enables structured evaluation of novel transcript detection. While the current implementation focuses on novel splice junctions, the method can be easily adapted to assess alternative transcription start sites (TSS), transcription termination sites (TTS), or transcript disambiguation by modifying the underlying GTF annotations. We demonstrated the utility of TUSCO in both human and mouse samples, with gene sets adaptable to tissue-specific or cross-tissue applications. This flexibility ensures that researchers working on tissues not directly included in the resource can still leverage the TUSCO framework. If adopted widely by the community, the approach could be extended to additional species where sufficient transcriptomic data exist to define a reliable set of single-isoform genes.

Despite its advantages, TUSCO also has limitations. It is inherently restricted to single-isoform loci and thus cannot evaluate transcript reconstruction accuracy within highly complex multi-isoform genes—a domain where synthetic spike-ins, with alternatively spliced genes, remain useful. However, our comparison with SIRVs shows that true positive (TP) rates were similar across both approaches, suggesting similar capacity for assessing the identification of the right junction chain. However, synthetic controls failed to capture sample-specific variation in pre-sequencing biases, as reflected by greater discrepancies in precision and false negative (FN) rates, which are better illuminated by the TUSCO gene-set. As sequencing error rates continue to drop, biases introduced during RNA extraction and library preparation are likely to become the dominant limiting factors. In this context, the endogenous nature of TUSCO genes offers a crucial advantage.

Another important consideration is the potential future discovery of alternative isoforms in currently annotated single-isoform genes. While the current TUSCO set is based on extensive sequencing evidence, continual updates and cross-referencing with evolving GENCODE annotations will be necessary to maintain its accuracy. Additionally, the framework has not yet been applied to conditions involving transcriptome divergence: miss-processed sample-specific transcripts that may accumulate, for example, in situations of cellular stress^48^. Although TUSCO has not been specifically applied to stress-related conditions, we propose it may be a useful tool for detecting transcriptome alterations under such scenarios, as rising false positive rates could reflect increased RNA processing errors associated with cellular stress.

In summary, TUSCO provides a powerful, biologically grounded framework to evaluate the performance of long-read transcriptomics across real-world conditions. By bridging the gap between synthetic controls and complex biological samples, it offers a more realistic benchmark to guide method development, protocol optimization, and experimental design.

## Methods

### TUSCO selection pipeline

TUSCO was constructed with a single, reproducible pipeline that integrates multi-source annotations, population splicing and TSS evidence, and broad expression compendia, and then enforces strict “no evidence for alternative isoforms” criteria. For human, we retrieved and harmonized GENCODE v48, RefSeq GRCh38.p14, and MANE v1.4 annotations together with an Ensembl BioMart Ensembl↔RefSeq/NCBI mapping; for mouse, we used GENCODE vM37 and RefSeq GRCm39 with the corresponding BioMart mapping (GENCODE, RefSeq/NCBI, MANE, Ensembl BioMart). After normalizing chromosome labels where needed, we identified genes that are single-isoform in each source and required exact identity of exon–intron structure and strand across all GTFs; only those cross-annotation matches were retained and propagated with synchronized mapping subsets.

We next screened for population-level alternative splicing and TSS inconsistencies. Splice evidence was derived from recount3 junction BEDs (canonical GT:AG sites for human; canonical-site filtering bypassed for mouse). For each multi-exon gene, we computed the mean coverage of annotated junctions, μ, and declared a novel junction as supporting an alternative isoform if it was fully intragenic, strand-consistent, and its coverage exceeded T = 0.03 × μ, with a minimum absolute cutoff of one supporting read; novel junctions shorter than 80 bp were ignored. TSS evidence was assessed with refTSS v4.1. In human, we required single-exon loci to pass both an exon-overlap check (exactly one refTSS interval must overlap the TSS exon and contain the transcript’s TSS coordinate) and a CAGE window check centered at the transcript TSS that removed loci with more than one peak in a ±300 bp window; multi-exon loci were also evaluated with the ±300 bp window. In mouse, TSS screening was restricted to single-exon genes using the exon-overlap rule without a CAGE window. For human only, we further removed genes flagged by IntroVerse_80 as having widespread novel isoform usage in population GTEx data (i.ed., novel-isoform evidence in more than 80% of samples).

Expression-based universality and tissue sets were then established using the resources and thresholds specified in the analysis configuration. For human, we used Bgee present/absent calls and RNA-Seq library metadata, retained calls of “gold quality”, “silver quality”, or “bronze quality”, and restricted analysis to tissues with at least 25,000 distinct expressed genes. Genes were called universal if present in at least 99% of retained tissues; anatomical entity IDs were mapped to tissue names and per-tissue gene sets were emitted using those IDs. For mouse, we used Bgee with the same minimum tissue gene coverage (25,000) and the same quality set, calling universal genes at a 90% prevalence threshold across retained tissues. To reinforce universality, we merged housekeeping genes from HRT Atlas (converted from transcript to gene identifiers via Ensembl REST) into the universal set for each species.

To ensure consistency at the population level beyond presence/absence, we applied an AlphaGenome-based universality screen to the post-expression gene set with species-specific thresholds. In human, genes were retained only if the median RPKM across mapped tissues was at least 2.0, if at least 99% of tissues showed RPKM exceeding 0.25, and if at least 99% of tissues had an unannotated-to-annotated splice ratio below 0.001. In mouse, the corresponding criteria were a median RPKM of at least 0.5, at least 90% of tissues above 0.01 RPKM, and at least 90% of tissues with splice ratio below 0.01. When reporting tissue-specific sets, an additional tissue-level AlphaGenome screen was applied using tissue RPKM and splice-ratio thresholds; for human we required RPKM > 6.0 and splice ratio < 0.001 in the tissue of interest, and for mouse RPKM > 1.5 and splice ratio < 0.001. A small, curated exclusion list was applied at the end to remove remaining edge cases. The pipeline writes synchronized mapping subsets alongside each GTF subset and records, for every removed gene, the first filter responsible of the removal. Final outputs include species-named GTFs (‘tusco_human.gtf’, ‘tusco_mouse.gtf’) and consolidated TUSCO tables with single-exon and multi-exon splits generated from the final selection.

### Sample Preparation, Library Construction, and Computational Pipeline Parameters

Long-read RNA-seq datasets used in this study were derived from multiple sources to enable a robust evaluation of transcriptome reconstruction using the TUSCO framework. Data were collected from both public repositories and our laboratory to provide diverse experimental conditions.

The LRGASP dataset comprises long-read sequencing data generated using Oxford Nanopore Technologies (ONT) and Pacific Biosciences (PacBio) platforms. These data, which include samples from human WTC11 induced pluripotent stem cells (iPSC) and mouse embryonic stem (ES) cells, were retrieved from the Synapse repository (https://www.synapse.org/Synapse:syn25007472/wiki/608702). The RNA degradation experiment data were obtained to assess the impact of sample quality on transcript recovery. FAST5 and FASTQ files for these experiments are available via the European Nucleotide Archive (ENA) under accession PRJEB53210, and additional FASTQ files from a post-mortem brain sample are accessible from the European Genome-Phenome Archive (EGA) under accession EGAS00001006542.

Mouse kidney dataset (PacBio Iso-Seq). Kidneys were dissected from five male C57BL/6J wild-type mice (3 months old). Tissues were homogenized using FastPrep, and total RNA was extracted with the Maxwell 16 LEV simplyRNA Purification Kit. cDNA synthesis and sample barcoding followed PacBio Iso-Seq recommendations using the NEBNext Single Cell/Low Input cDNA Synthesis & Amplification kit. Five barcoded Iso-Seq libraries (K31–K35) were prepared with the Iso-Seq SMRTbell prep kit 3.0 and sequenced on a PacBio Sequel IIe. This dataset will be made publicly available upon publication.

Mouse-specific laboratory procedures. RNA quantity and integrity were assessed prior to library preparation (e.g., Agilent Bioanalyzer to obtain RIN values). No direct-RNA or ONT libraries were generated for this mouse dataset. Library preparation strictly followed the PacBio Iso-Seq workflow: full-length cDNA amplification and barcoding, SMRTbell construction with the Iso-Seq SMRTbell prep kit 3.0, and sequencing on the Sequel IIe platform. Run and enzyme/incubation parameters followed the manufacturer’s recommendations for Iso-Seq libraries. Lexogen SIRV-Set 1 spike-ins were added at 3%. Specifically, we added E0 mix to brain samples and E1 mix to kidney samples

For all dataset, alignment of reads to the reference genome was conducted using minimap2 (version 2.0) via a custom script. The reference genome mm39, provided in FASTA format (e.g., ref_chr_primary_assembly.fasta), was pre-indexed using minimap2, and alignments were performed with parameters specifically tuned for long-read RNA-seq data. Key settings included the use of the -ax splice option for spliced alignment, --secondary=no to suppress secondary alignments, and -C5 to penalize ambiguous mappings. The flag -uf was used control correct splice site detection, and the maximum intron length was set to 2,000,000 nucleotides via the -G 2000000 parameter. Code is provided as Supplementary File S2.

For transcript quality control and classification, the SQANTI3 pipeline (version 5.0) was applied. In general, the pipeline uses reference annotation, genome, and CAGE peak data specific to the organism under study. The workflow is divided into three major stages: a quality control (QC) stage using sqanti3_qc.py, a filtering stage using sqanti3_filter.py that applies criteria defined in a default JSON configuration file to remove low-confidence transcript models, and a rescue stage using sqanti3_rescue.py to recover valid transcripts that may have been overly filtered^16^.

We ran the pipeline with common options such as --skipORF to bypass open reading frame prediction and provided Illumina short-read coverage files, obtained by STAR mapping, of the same samples to support splice junction validation.

### Transcriptome Reconstruction with FLAIR

We reconstructed transcriptomes for three datasets —(i) the LRGASP project (Oxford Nanopore [MinION R10.4.1] and PacBio Sequel II), (ii) the RNA degradation dataset (Oxford Nanopore direct RNA [GridION R9.4.1]), and (iii) the replicated Mouse Brain & Kidney dataset (PacBio Sequel IIe and Oxford Nanopore cDNA [R9.4.1])—using FLAIR v2.0^49^. Briefly, raw long-read sequencing data were first converted to FASTQ format. Reads were then aligned to the reference genome using FLAIR’s align functionality, which internally wraps minimap2 to produce BED-formatted alignments. FLAIR was run without Illumina short-read correction/filtering. Alignment files were refined with the correct module to adjust splice junctions and transcript boundaries based on an external reference GTF annotation, thereby rectifying mis-mappings often attributable to error-prone ONT signals.

### TUSCO metrics calculation

We curated a single-isoform GTF reference (“TUSCO reference set”) comprising genes with no evidence of alternative splicing. Using the SQANTI3 classification file, we restricted evaluation to reconstructed transcripts that map to TUSCO genes, matched by the predominant identifier present in the classification (Ensembl, RefSeq, or gene symbol). We labeled as true positives (TP) those reference_match transcripts that reproduce the TUSCO exon-intron structure, including qualifying mono-exon calls when the reference has one exon and both TSS and TTS are within +/-50 bp; partial true positives (PTP) were full-splice_match or incomplete-splice_match transcripts that deviate at the 5’ or 3’ ends; false positives (FP) were transcripts within TUSCO genes labeled novel_in_catalog, novel_not_in_catalog, genic, or fusion; and false negatives (FN) were TUSCO genes with no observed transcript in any structural category. Let G be the number of TUSCO genes, N the number of observed transcripts mapping to TUSCO genes, TP the number of TP transcripts, PTP the number of PTP transcripts, FP the number of FP transcripts, TP_g the number of TUSCO genes with at least one TP transcript, D the number of TUSCO genes detected by either TP or PTP, and T_fsm+ism the total number of full-splice_match plus incomplete-splice_match transcripts. From these quantities we calculated: Sn = 100 * TP_g / G; nrPre = 100 * TP / N; rPre = 100 * (TP + PTP) / N; PDR = 100 * D / G; FDR = 100 * (N - TP) / N; FDeR = 100 * FP / N; redundancy = T_fsm+ism / D.

### Intersection mode across replicates

For replicate analyses, we computed TUSCO metrics on a k-way intersection call set that retains only transcript structures reconstructed in all of the k selected replicates. Within each replicate, SQANTI3- classified transcripts were first assigned to TUSCO genes and collapsed by structure (identical junction chain; TSS/TTS considered equivalent within ±50 bp). A structure entered the intersection if an equivalent collapsed transcript appeared in every replicate; one representative was then chosen by (i) minimum summed 5′/3′ end deviation from the TUSCO model, (ii) higher read support, (iii) longer length. Correctness labels (TP, PTP, FP) and all downstream metrics (Sn, nrPre, rPre, PDR, FDR, FDeR, redundancy) were computed exclusively on this intersection set; FN were TUSCO genes lacking TP or PTP in the intersection. This conservative strategy emphasizes reproducibility, sharply reducing spurious calls while preserving consistently detected transcripts.

### Data availability and code availability

Publicly available long-read RNA-seq datasets used in this study include the LRGASP data for human WTC11 iPSC and mouse ES cells, accessible at the Synapse repository under Synapse: syn25007472/wiki/608702. The RNA degradation datasets are deposited at the European Nucleotide Archive (ENA) under accession PRJEB53210, with supplementary FASTQ data for post-mortem brain samples available from the European Genome-Phenome Archive (EGA) under accession EGAS00001006542. The Replicated Mouse Brain and Kidney dataset, generated in-house, has been submitted in its entirety to the ENA under accession PRJEB94912 (secondary accession ERP177675, “Sequencing of mice kidneys using IsoSeq”). The dataset will be released publicly upon publication.

For benchmarking, the universal and tissue-specific TUSCO gene sets for human, and mouse can be freely obtained at https://github.com/ConesaLab/SQANTI3 and at the dedicated TUSCO portal https://TUSCO.uv.es.

All scripts used to generate the figures in this study and to implement the TUSCO gene selector are available at https://github.com/TianYuan-Liu/tusco-paper.

## Supplementary Figure

**Figure S1:**
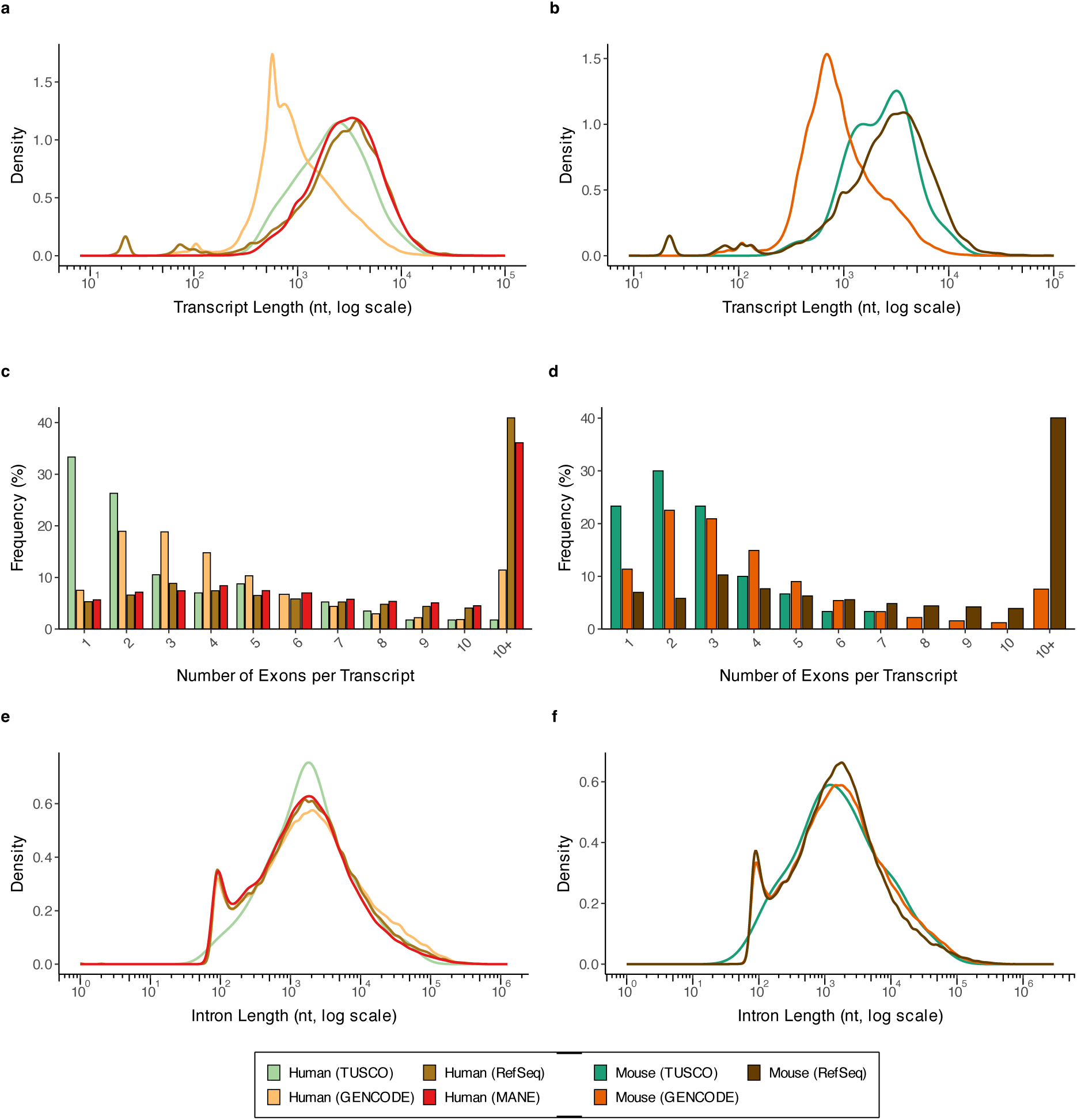
Comparative analysis of transcript features in TUSCO versus reference annotations for human and mouse. (a,b) Density plots of transcript lengths (log10 scale) in human (a) and mouse (b) show the distribution of TUSCO transcripts (light green in human, darker green in mouse) relative to RefSeq (golden brown in human, dark brown in mouse), GENCODE (light orange in human, darker orange in mouse), and MANE (red, human only). In both species, TUSCO transcripts tend to have length profiles comparable to or slightly longer than major annotation sets. (c,d) Frequency histograms of exon count per transcript in human (c) and mouse (d), using the same color scheme: TUSCO (light/darker green), RefSeq (golden/dark brown), GENCODE (light/darker orange), and MANE (red, human only). Although TUSCO is enriched for single-exon genes, it also includes multi-exon transcripts spanning a broad exon count range. (e,f) Density plots of intron lengths (log10 scale) for human (e) and mouse (f) similarly demonstrate that TUSCO (light/darker green) largely overlaps with RefSeq, GENCODE, and MANE in intron architecture. Collectively, these comparisons confirm that TUSCO transcripts overlap with naturally occurring size ranges and gene structures found in standard reference annotations, making TUSCO an internally consistent benchmark for long-read transcriptomic quality assessments.

**Figure S2.**
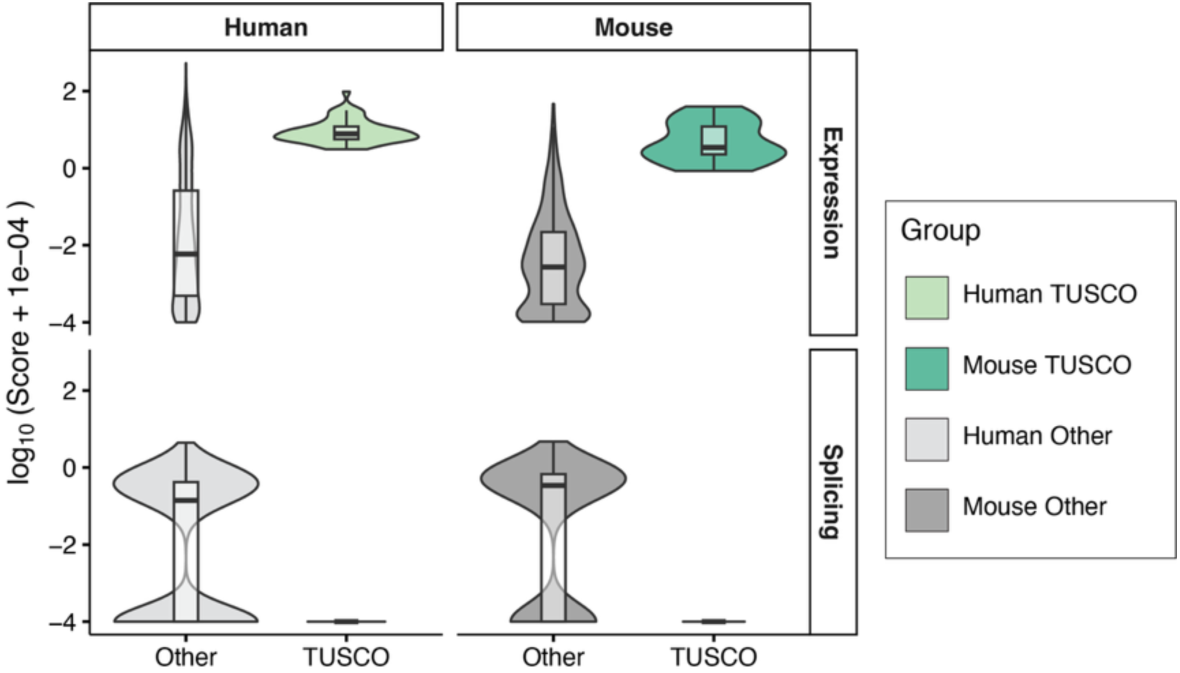
TUSCO genes are highly expressed and display minimal novel splicing signal. Violin– box plots compare AlphaGenome-derived scores for TUSCO genes versus other single-isoform genes in human and mouse. The top row shows the Expression score (gene-level tissue-median RPKM); the bottom row shows the Splicing score (maximum splicing score/ratio per gene). Values are plotted as log₁₀ (score + 1e−4). Across both species, TUSCO genes exhibit substantially higher expression and lower novel splicing score ratios than other single-isoform genes, consistent with their selection as widely expressed, splicing-invariant controls. Box centers mark medians; boxes span the interquartile range; whiskers extend 1.5× IQR. Colors: Human TUSCO (light green), Mouse TUSCO (teal), Human Other (light gray), Mouse Other (dark gray).

**Figure S3.**
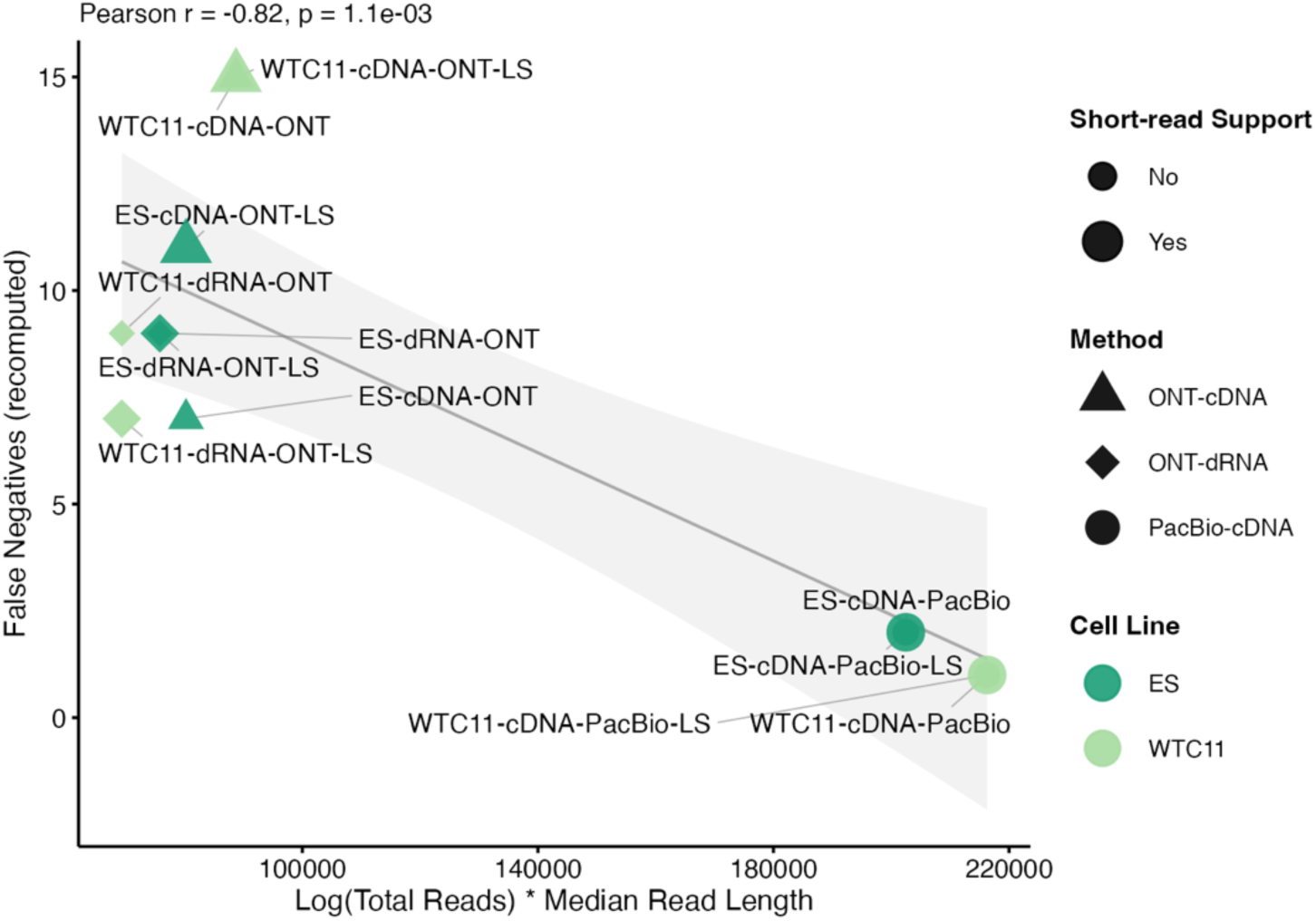
False negatives decrease with effective sequencing yield. Scatter plot showing the number of recomputed TUSCO false negatives (y-axis) versus a composite metric, Log (Total Reads) × Median Read Length (x-axis), across LRGASP datasets. Point shape encodes method (triangle: ONT-cDNA; diamond: ONT-dRNA; circle: PacBio-cDNA), color indicates cell line (ES, WTC11), and point size denotes short-read support (larger = yes). Labels include dataset names; “-LS” marks libraries with length selection. A linear fit with 95% confidence band (gray) reveals a strong negative correlation (Pearson r = −0.82, p = 1.1 × 10⁻³), indicating that higher effective depth and longer reads substantially reduce false negatives, consistent with technical rather than biological causes.

**Figure S4:**
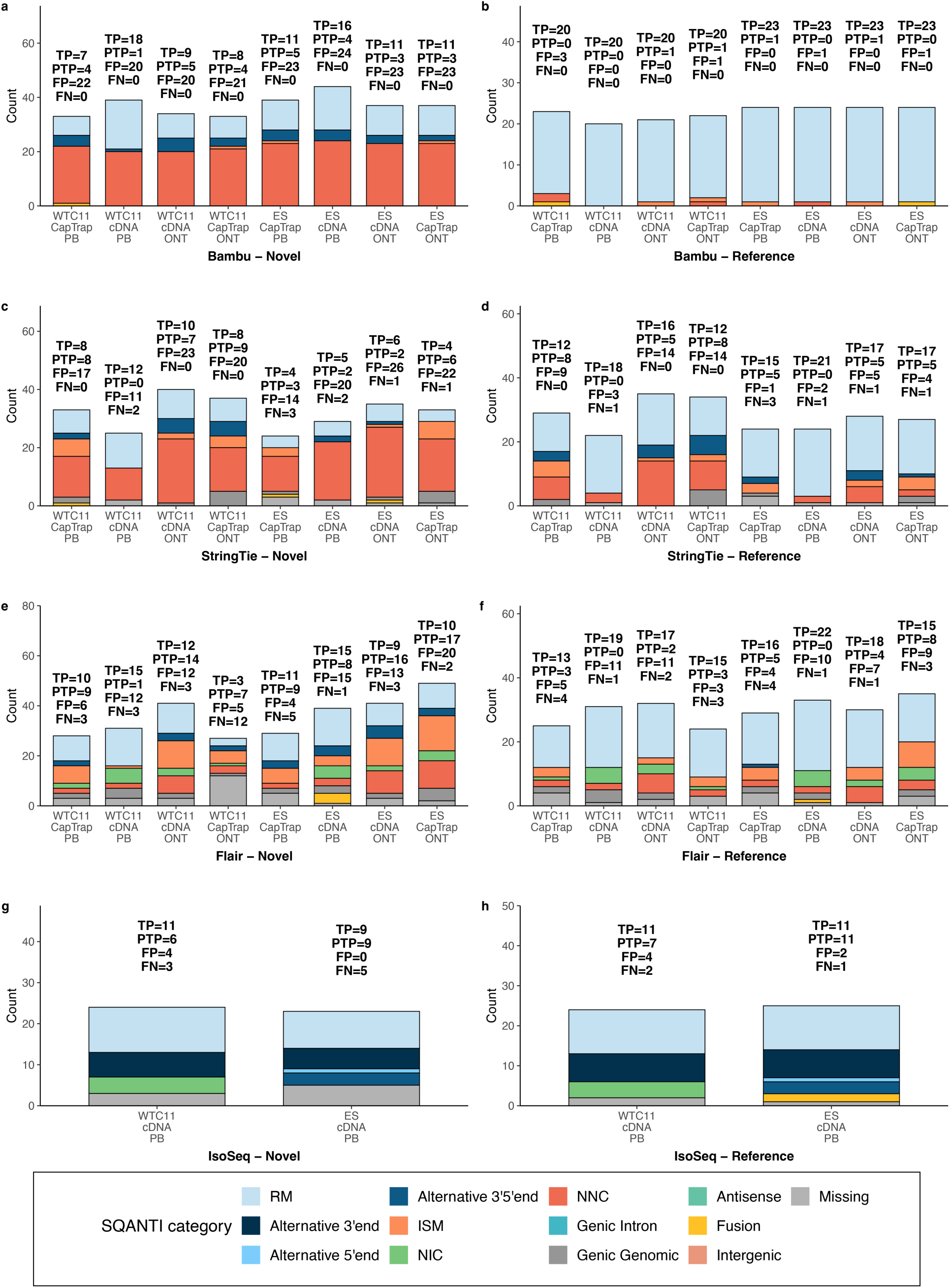
Performance of long-read transcriptome reconstruction pipelines on the TUSCO benchmark in discovery unannotated genes. Panels a–h (a, Bambu-Novel; b, Bambu-Reference; c, StringTie-Novel; d, StringTie-Reference; e, Flair-Novel; f, Flair-Reference; g, IsoSeq-Novel; h, IsoSeq- Reference) show stacked-bar summaries for each human (wtc11-*) and mouse (es-*) sample: total bar height equals the number of multi-exon TUSCO genes assessed, colours indicate SQANTI3 structural categories (legend). Turquoise RM segments correspond to full splice matches with identical transcription start and termination and are scored as true positives (TP); alternative 3′, 5′ or 3′/5′ ends plus ISM represent partial true positives (PTP); NIC, NNC and other genic/intergenic classes constitute false positives (FP); grey “Missing” portions are false negatives (FN). Numeric overlays list TP, PTP, FP and FN for each bar. In the Novel regime the authentic TUSCO isoform was replaced by a synthetic splice-junction variant (TUSCO-novel), forcing pipelines to rediscover the true isoform in the presence of a misleading annotation, whereas the Reference regime uses the unedited annotation to measure classical recall. The figure therefore captures, in a single glance, both absolute recovery of true isoforms and the mis-classification spectrum that arises when annotation incompleteness mimics real-world transcript discovery scenarios.

**Figure S5.**
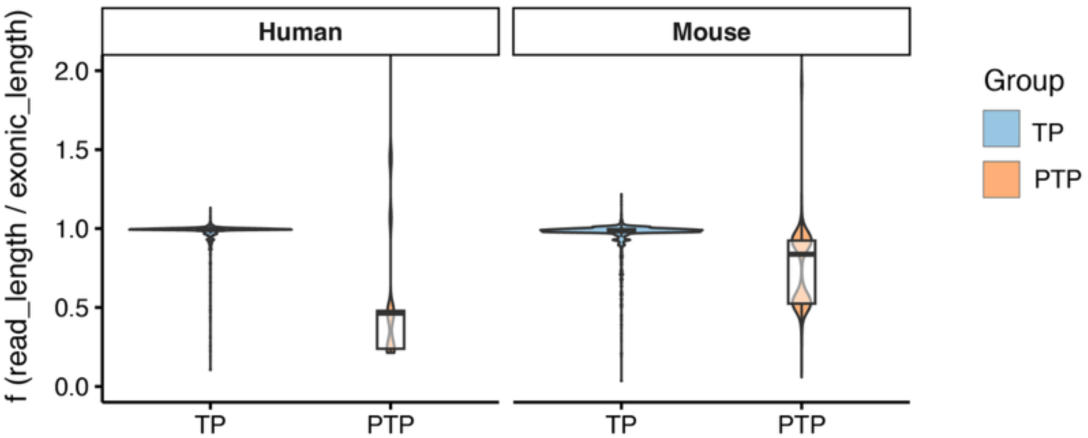
Per-read coverage of TUSCO transcripts in LRGASP Iso-Seq cDNA data. Violin-plus- boxplots showing the distribution of per-read coverage f = read_length/exonic_length for TUSCO-mapped reads, faceted by species (Human WTC11, Mouse ES). Reads are labeled by SQANTI as TP (full-splice match; reference_match) or PTP (incomplete/partial splice match). Only multi-exon TUSCO transcripts are included; reference- and novel-evaluation classifications are combined; no TSS-based filtering; outliers are retained; y-axis limited to 0–2.0. TPs cluster near full-length, consistent with accurate reconstruction (mean 95.6% of transcript length; n=10,921), while PTPs are shorter, consistent with partial transcripts due to sample and library biases (mean 75.1%; n=2,731). By species: Human—TP 95.1% (n=6,716), PTP 67.5% (n=176); Mouse—TP 96.5%

**Figure S6:**
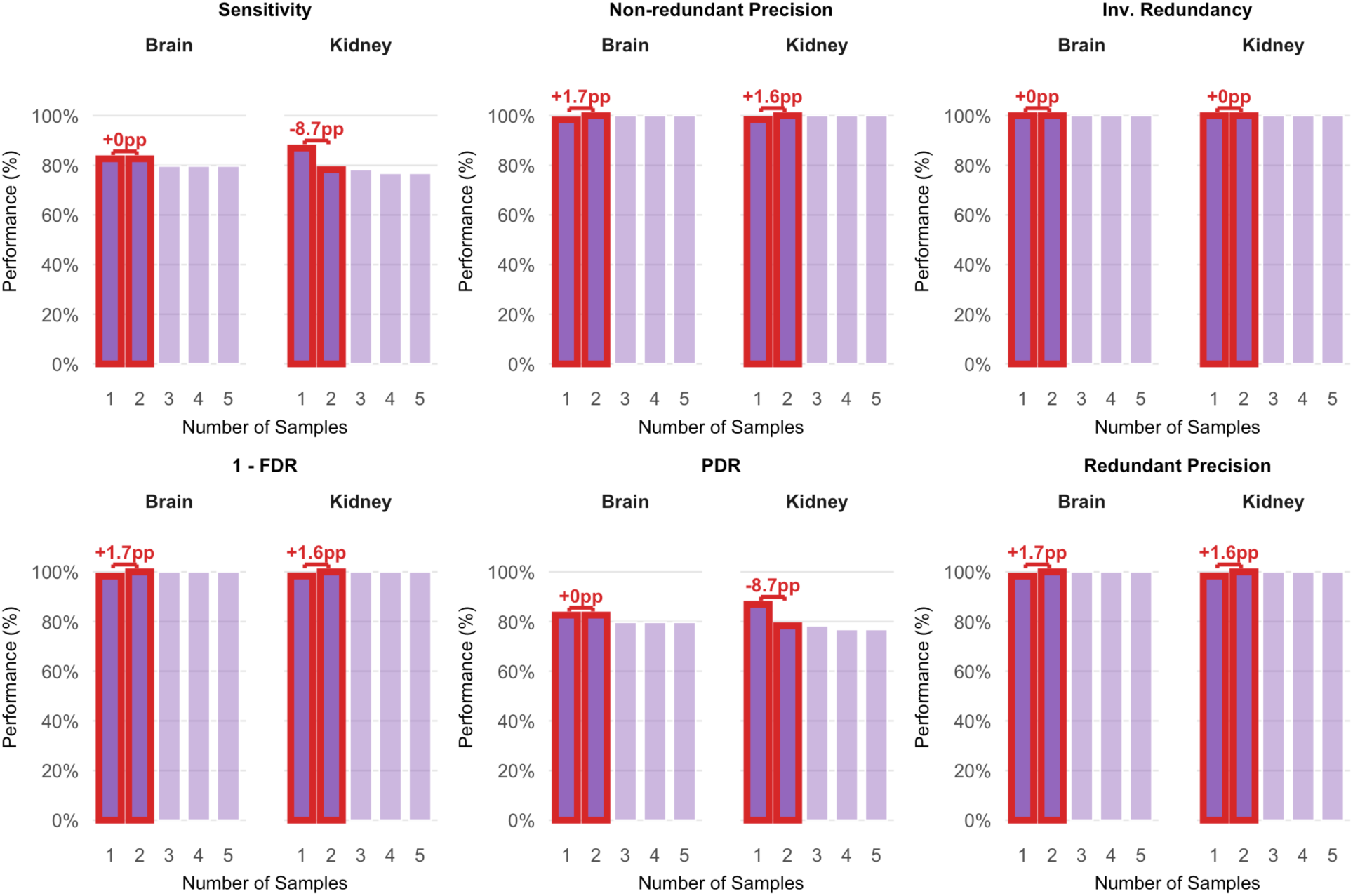
SIRV spike-in transcript reconstruction improves modestly but consistently with additional replicates. Purple bars show performance on the SIRV controls in mouse brain (left columns of each panel) and kidney (right columns) as replicate depth increases from one to five libraries. Performance is scored only for transcripts reproducibly detected across all replicates. Red annotations indicate the percentage-point change upon adding a second replicate (1 → 2 samples). (a) Sensitivity remains unchanged in brain (+0 pp) but falls modestly in kidney (–8.7 pp), reflecting the stricter consensus requirement. (b) Non-redundant precision rises by +1.7 pp (brain) and +1.6 pp (kidney), demonstrating improved accuracy in calling unique isoforms. (c) Inverse redundancy reaches 100 % after a single replicate and shows no further change (+0 pp). (d) 1 – FDR increases by +1.7 pp (brain) and +1.6 pp (kidney), indicating reduction of false positives. (e) Positive detection rate (PDR) is stable in brain (+0 pp) but decreases in kidney (–8.7 pp) under the higher consensus threshold. (f) Redundant precision improves by +1.7 pp in brain and +1.6 pp in kidney, mirroring the gains in non-redundant precision. Beyond two replicates, all metrics plateau (< 1 pp change), indicating diminishing returns.

## Supplementary table

**Table S1.** Cosine similarity (cosim) scores across different pipelines for mouse embryonic stem cells (ES) and human WTC11 cells. Cosine similarity values close to 1 indicate high agreement. LS denotes pipelines that integrate long and short reads.

|  | WTC11 | ES |
| --- | --- | --- |
| cDNA ONT | 0.974 | 0.991 |
| cDNA ONT LS | 0.975 | 0.988 |
| cDNA PacBio | 0.999 | 0.999 |
| cDNA PacBio LS | 0.999 | 0.999 |
| dRNA ONT | 0.996 | 0.993 |
| dRNA ONT LS | 0.998 | 0.993 |

**Table S2.** Read counts for PacBio Iso-Seq samples from mouse kidney (K31–K35) and brain (B31–B35). Each value represents the total number of reads mapped in the corresponding sample, as determined by samtools. The “Tissue” column indicates the biological source of each sample.

|  | Read Count | Tissue |
| --- | --- | --- |
| K31 | 4,593,279 | Kidney |
| K32 | 5,136,502 | Kidney |
| K33 | 3,902,779 | Kidney |
| K34 | 5,436,609 | Kidney |
| K35 | 5,096,149 | Kidney |
| B31 | 3,684,709 | Brain |
| B32 | 3,684,709 | Brain |
| B33 | 3,854,778 | Brain |
| B34 | 4,413,078 | Brain |
| B35 | 3,888,139 | Brain |

